# Inbreeding and resultant homozygosity across key inflammation and DNA repair genes linked to chlamydial infection in New South Wales koalas

**DOI:** 10.1101/2025.07.22.666211

**Authors:** Elspeth A. McLennan, Larry Vogelnest, Luke W. Silver, Belinda R. Wright, Andrea Casteriano, Zhiliang Chen, Valentina S. A. Mella, Damien P. Higgins, Mark B. Krockenberger, Katherine Belov, Tim S. Jessop, Carolyn J. Hogg

**Author notes:** Corresponding author: Professor Carolyn Hogg |.

## Abstract

Inbreeding and resultant homozygosity can reduce genetic diversity and increase disease susceptibility. Koalas (*Phascolarctos cinereus*) are one species suffering genomic diversity loss and inbreeding and concurrent significant disease pressure (particularly chlamydiosis). Using 259 whole genomes with a pathogen sampling regime we identify potential links between inbreeding, genome-wide variation and chlamydial infection. We found a general trend of reduced genomic diversity and increased inbreeding from north to south across six sites in New South Wales. A genome-wide association study of 153 individuals from sites with known *Chlamydia pecorum* presence were used to investigate the potential relationship between inbreeding and infection. *Chlamydia* positive individuals (average FH = 0.026) were significantly more inbred than *Chlamydia* negative individuals (average FH= −0.0051) (*t* = - 2.31, df = 151, p-value = 0.022). We identified several genes involved in host-pathogen interactions and DNA mismatch repair within in runs of homozygosity that were unique to *Chlamydia* positive individuals. Interestingly, populations considered putatively *Chlamydia*-free had similar allele frequencies across candidate loci as *Chlamydia* positive individuals. Combined with gene flow analyses, this result suggests that isolation may have protected these populations more than harbouring alleles conferring infection resilience and supports the concept that disease should be carefully considered in any conservation measures that increase connectivity or translocations. Our genome-wide approach has identified several avenues for investigations into the pathogenesis of *Chlamydia* infection and chlamydiosis. We showcase the value of high-quality re-sequenced genomes for understanding the implications of inbreeding, genomic diversity loss, and infection susceptibility, all universal problems for threatened species.

## Introduction

For a range of global species, inbreeding is accumulating as a result of fragmented populations and/or small population sizes resulting in reduced genomic diversity, and a reduction of adaptive potential of that population/species to respond to either current or future events (Shaw et al., 2025). Inbreeding, and subsequent exposure of deleterious alleles, is also predicted to increase disease susceptibility (Hedrick & Garcia-Dorado, 2016). For example, female European badgers (*Meles meles*) suffered exacerbated progression of bovine tuberculosis with higher levels of inbreeding (Benton et al., 2018). Similarly, inbred horse nettle plants (*Solanum carolinense*) were more susceptible to powdery mildew (*Oidium neolycopersici*) and suffered more rapid onset of infection compared to their outbred counterparts (Kariyat et al., 2012). Advances in sequencing technology have enabled more accurate assessments of genome-wide inbreeding via detection of runs of homozygosity (ROH) (Ceballos et al., 2018). ROH are large homozygous sections of the genome identical by descent from a common ancestor (Ceballos et al., 2018). Longer ROH represent recent inbreeding, whereas shorter ROH are indicative of historic inbreeding that has been broken apart by recombination with non-relatives (Ceballos et al., 2018; Curik et al., 2014). ROH analyses can be used to detect homozygosity across key genes associated with responses to infectious diseases and other fitness traits (Colpitts et al., 2022; Silver et al., 2025; Wilder et al., 2024). Understanding the link between low diversity as a result of inbreeding across key response genes and disease susceptibility may help pinpoint potentially vulnerable individuals or populations to known disease threats.

Infectious diseases of wildlife are increasingly being recognised as a significant problem in conservation (Barroso et al., 2021). Key examples include frogs and chytrid fungus (Fisher & Garner, 2020), bats and white-nose syndrome (Cheng et al., 2021), and Tasmanian devils (*Sarcophilus harrisii*) and devil facial tumour disease (Clarke et al., 2019), all of which have caused substantial population declines. Infectious agents are a part of ecosystem dynamics, but the current anthropogenic pressure on wildlife and their habitats through fragmentation and loss (Crooks et al., 2017) has resulted in increased exposure to novel pathogens and zoonotic spillover (Glidden et al., 2021). Not only are species being exposed to a wider range of pathogens, but their adaptive potential to effectively co-evolve is being eroded through increased isolation, inbreeding, and loss of genetic diversity (Glidden et al., 2021; Shaw et al., 2025).

The koala (*Phascolarctos cinereus*) is a species facing a range of threats including habitat loss and fragmentation (McAlpine et al., 2015), leading to an increase in inbreeding and loss of genomic diversity (McLennan et al., 2025a), as well as facing strong disease pressures (Vitali et al., 2023). Genomic diversity and inbreeding is not uniform across the species’ range, but generally decreases and increases respectively on a north to south cline, with the exception of isolated sites along the east coast (McLennan et al., 2025a). This study also showed there to be limited contemporary gene flow between the sites studied, further increasing the likelihood of ongoing reductions in genomic diversity (McLennan et al., 2025a). Chlamydiosis, caused by the bacterium *Chlamydia pecorum*, is present across many koala populations and has caused significant population declines in some regions (Quigley & Timms, 2020; Rhodes et al., 2011). However, prevalence of infection and disease severity vary across the koala’s range (Quigley & Timms, 2020). A previous study found no association between inbreeding and chlamydial infection in a single population in Queensland, using 5,000 single nucleotide polymorphisms predisposed to neutral regions of the genome (Cristescu et al., 2022). Until now, no multi-population, whole-genome assessments have been made to investigate associations between inbreeding, potential candidate loci/genes and chlamydial disease for multiple koala populations.

Chlamydiosis is primarily a sexually transmitted disease that can also be transferred vertically from females to joeys (Jackson et al., 1999; Nyari et al., 2017). *C. pecorum* is an obligate intracellular pathogen that targets mucosal surfaces, including conjunctiva and urogenital tract, and predominantly invades epithelial cells (Kayesh et al., 2024). Though it likely persists and traffics between anatomical sites *via* histiocytic cells, such as macrophages, Langerhans cells or dendritic cells, as it does in other species (Burach et al., 2014; Higgins et al., 2005). As *C. pecorum* grows and replicates within cells it disrupts the normal cellular functions (Kayesh et al., 2024). Local and systemic inflammatory responses are initiated and directed by infected and neighbouring epithelial cells, histiocytic antigen presenting cells and other sentinel cells, and modulated and effected by finely balanced pathways including phagocytosis, cytokine activation and T-cell responses (Pagliarani et al., 2024; Phillips et al., 2024). Chlamydial disease in koalas is associated with chronic inflammation of the reproductive, urinary and ocular regions and includes but is not limited to ovarian bursal cysts, male and female reproductive tract inflammation, keratoconjunctivitis, pyelonephritis, urethritis and cystitis. Impacts can therefore include infertility, blindness, ill-thrift and death and, with incontinence, the characteristic “wet bottom” or brown staining of the fur around the cloaca (Hemsley & Canfield, 1997; Higgins et al., 2005; Hulse et al., 2022).

As the immune system is responsible for the initial response to infection and development of disease, research into the susceptibility of koalas to *Chlamydia* has so far focused on immune genes. Associations between various genotypes of the major histocompatibility complex (MHC) class I and II genes, have been associated with various aspects of chlamydial infections and disease progression (Kidd et al., 2024; Lau et al., 2014; Lau et al., 2013; Quigley et al., 2018; Silver et al., 2022). Some of the diverse humoral and cellular pathways associated with chronic infection and disease have been described in koalas (Higgins et al., 2005; Quigley & Timms, 2021). However, our understanding of factors that affect host-pathogen relationships early in chlamydial infection of koalas, when infection is established and subsequent immune pathways are determined, is less clear. The wide range of host cell and inflammatory pathways involved and the diversity of metabolic, stress, and reproductive pathways that can influence them (Downs et al., 2014) suggests the benefit of a broad approach to examining the genetic architecture that underpins these processes and its relationship to chlamydial infection and persistence. Here, we employed a genome-wide approach to determine whether inbreeding and homozygosity across any gene family were linked to *Chlamydia* infection in koalas, as well as detecting candidate loci through a traditional case-control approach.

## Materials and Methods

### Study species & sampling sites

Although a comparatively well-studied species, there remain significant knowledge gaps pertaining to the current ecology, genomics and diet of koalas across New South Wales (NSW) all of which may be important factors in disease dynamics. Without such valuable information, the impact from threatening processes cannot be accurately assessed, nor the potential population responses be predicted. Such knowledge is critical for determining the best management actions to safeguard koala populations and maximise future viability. As such, the NSW Government (Department of Climate Change, Energy, Environment, and Water), in collaboration with the NSW National Parks and Wildlife Service, Taronga Conservation Society Australia (Taronga), The University of Sydney, The Australian National University, and the University of Western Sydney, launched the NSW Koala Sentinel Program that aimed to gain detailed baseline information, including reproduction, mortality, dispersal, genomic diversity, population connectivity, inbreeding, diet, microbiome, and disease prevalence and transmission, across six key locations, or sentinel sites, for NSW koalas.

The sentinel sites are located in the Richmond Range National Park (hereafter Richmond Range; 28.807°S, 152.742°E), Port Macquarie Conservation Areas (hereafter Port Macquarie; 31.435° S, 152.900° E), Kanangra-Boyd National Park (hereafter Kanangra-Boyd; 33.832° S, 149.999° E), Woronora Plateau Conservation Areas (hereafter Woronora Plateau; 34.037° S, 151.023° E), the Southern Tablelands Conservation Areas (hereafter Southern Tablelands; 34.885° S, 149.472° E), and the Narrandera Common and Conservation Areas (hereafter Narrandera; 34.748° S, 146.550° E) (Figure 1a). These sites are geographically and climatically representative of koala distributions in NSW, perceived important koala strongholds with large populations, often with information available from prior research projects, and were chosen considering local stakeholders including First Nations communities and the cultural significance of koalas in those areas (DCCEEW, 2024).

**Figure 1.**
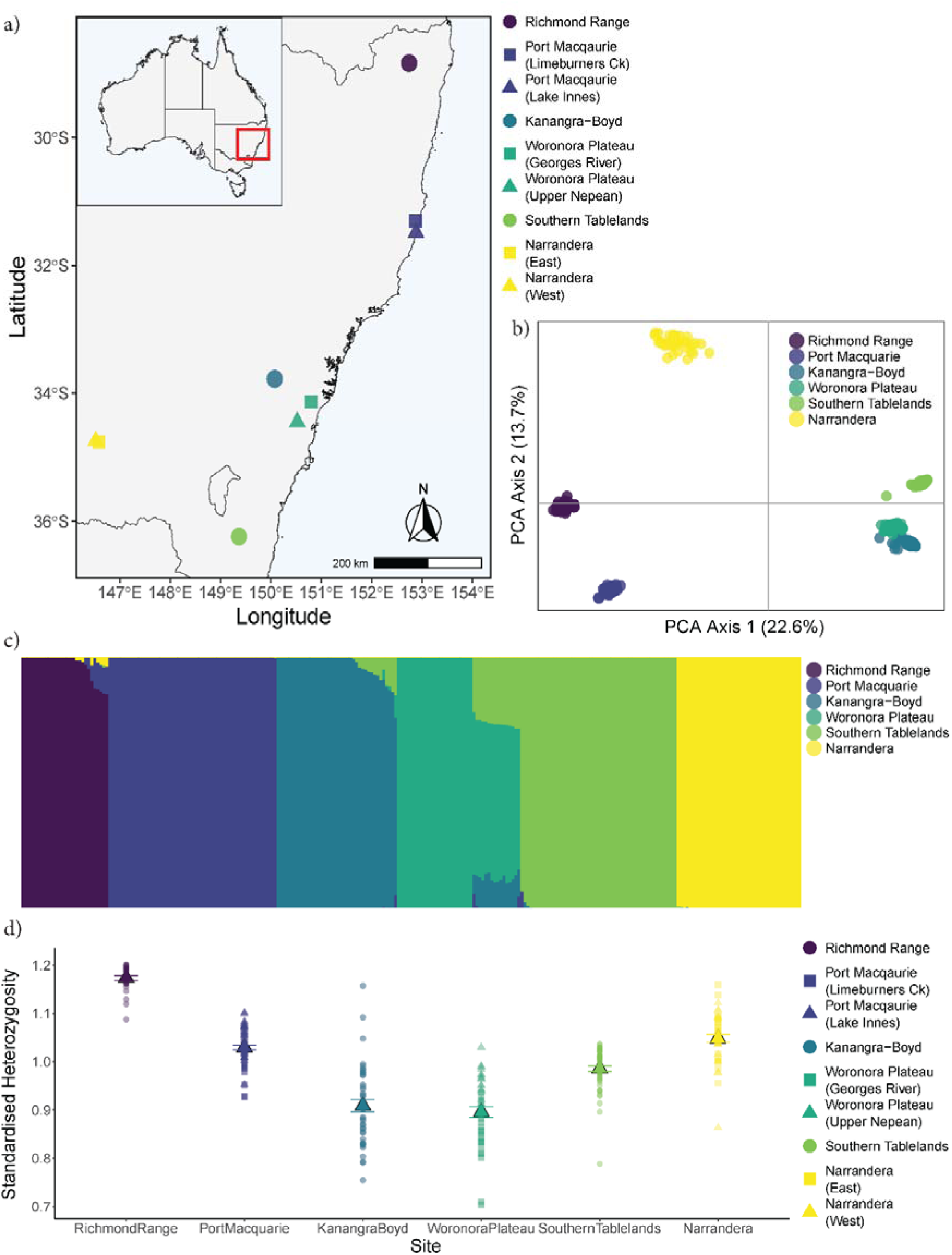
a) Map of New South Wales Sentinel Program sites with different symbols representing the subsites where relevant and inset map of Australia for geographic context, b) Principal component analysis of the six sentinel sites, c) *fastSTRUCTURE* plot showing the genetic ancestry assignment of each individual assigned across genetic clusters which are represented by sampling sites, d) standardised heterozygosity of each sampling site, the average represented by triangles and error bars representing standard error, scatter points represent the spread of the data and symbols relate to site/subsite where relevant.

Within three sites, Port Macquarie, Woronora Plateau and Narrandera, there was sufficient geographic spread in the sampling area and/or potential large barriers to gene flow, including townships, for two separate sampling locations to be included to capture koalas across the broader range of the area. The subsites include Limeburners Creek (31.307° S, 152.867° E) and Lake Innes (31.491° S, 152.818° E) within Port Macquarie, Georges River (33.964° S, 150.981° E) and the Upper Nepean (34.418° S, 150.617° E) within Woronora Plateau, and East (34.762° S, 146.586° E) and West (34.744° S, 146.522° E) Narrandera, either side of the township. Summary population genomic statistics (described below) were calculated at both the site and subsite level where appropriate.

### Sample collection

Between November 2023 and November 2024, 300 koalas were sampled during routine koala sentinel monitoring trips. Here, koalas were detected by drone, then captured using ‘guide rope and flag’ method routinely employed for koala capture (Madani et al., 2020), and various samples for disease and genetic analysis were collected during veterinary examination. For health assessments and sample collection, koalas were anaesthetised using a combination of alfaxalone (2 mg/kg) and medetomidine (0.05 mg/kg) administered intramuscularly, which lasted approximately 45-50 mins. A range of samples and morphometric data were collected. Relevant to this study, ear tissue biopsies were collected and stored in 70% ethanol at −80°C. For the detection of *Chlamydia* spp. bacteria, urogenital and ocular swabs were collected using a Copan Minitip FLOQSwab® (Copan, Brescia, Italy) (see Supplementary Methods for full anaesthesia and sampling regime).

Koala DNA for whole genome sequencing was extracted from tissue samples using the MagAttract HMW DNA kit (Qiagen, Germany). DNA repair was performed using the FFPE DNA repair protocol (New England Biosciences, USA) to improve DNA quality and ensure consistency in sample sequence coverage as per Hogg et al. (2023). DNA concentration was quantified using a Qubit 4.0 Fluorometer (ThermoFisher Scientific, USA) and quality assessed on a 1% agarose gel run for 30 minutes at 90V.

For detection of *Chlamydia* shedding, genomic DNA was extracted from the urogenital and ocular swabs using the MagMAX CORE Nucleic Acid Extraction kit (ThermoFisher, Scientific) and a KingFisher™ Flex purification system. DNA was eluted to 100 µL final volume. The beta-actin mRNA (housekeeping) gene, chlamydial genus 23S rRNA and *C. pecorum ompB* gene were targeted in a multiplexed qPCR assay using primers and probes previously described (Hulse et al., 2018). PCR reactions were made up to a final volume of 20 µL including 10 µL of SensiFAST Probe No-ROX (x2), 400 nM of each primer pair, 200 nM of each probe, and 2 µL of DNA template. PCR conditions included an initial denaturation step at 98°C for two minutes, followed by 40 cycles at 98°C for 15 seconds denaturation, followed by annealing/extension at 58°C for 40 seconds. Samples were run at full concentration and in duplicate. The limit of detection for this assay is 86 target copies per reaction (95% CI), determined using Probit regression analysis. Individuals were considered *Chlamydia* positive in the context of this study if one or both (ocular and urogenital) swabs returned a positive result for *Chlamydia* shedding.

*Chlamydia* shedding was considered present, i.e., the sample was positive, if amplification occurred for both the 23S and *ompB* targets and negative if amplification occurred at neither target. Samples with low or weak 23S amplification but no *C. pecorum* specific amplification were further tested using a different set of primers which target a different portion of the 23S sequence, to confirm the presence of *Chlamydia*. This confirmatory qPCR assay was made up to a final volume of 20 µL including 10 µL of SsoAdvanced Universal SYBR Green Supermix (x2), 250 nM of each primer (Ehricht et al., 2006) and 2 µL of DNA template. PCR conditions included an initial denaturation step at 98°C for two minutes, followed by 40 cycles at 98°C for 15 seconds denaturation, followed by annealing/extension at 60°C for 30 seconds, with melt curve analysis from 55 - 95°C at 0.5°C increments for 5 seconds. Amplification products with melt temperatures within the expected range were considered positive for *Chlamydia*.

### Whole genome re-sequencing

Whole genome re-sequencing was performed on 259 samples from across the six Sentinel sites (*N* = 29 Richmond Range; *N* = 56 Port Macquarie, *N* = 40 Kanangra-Boyd, *N* = 41 Woronora Plateau, *N* = 52 Southern Tablelands, and *N* = 41 Narrandera). Sequencing was performed at the Ramaciotti Centre for Genomics (The University of New South Wales) with sample libraries prepared using the Illumina DNA PCR-free prep and sequencing performed using the NovaSeq X Plus platform (Illumina, USA) with 150 bp pair-end reads.

Raw sequence reads were aligned to the koala reference genome (Johnson et al., 2018), scaffolded using Dovetail HiC (GCA_003287225.2_phaCin_HiC), using the DRAGEN Germline Pipeline (v4.3.6; Illumina Connected Analytics). Joint genotyping was performed across all samples (gVCF files) in batches of 100 using the DRAGEN Joint Pedigree Calling Pipeline (v4.3.6; Illumina Connected Analytics) resulting in three separate VCF files. VCF files were merged using PLINK v1.9 (Chang et al., 2015) to create an input file with joint genotyping across all 259 koala samples.

### Population genomics

#### 1. Population structure

For assessments of population structure and basic population summary statistics, single nucleotide polymorphisms (SNPs) were filtered using VCFtools v0.1.16 (Danecek et al., 2011) based on a call rate of 0.9, minor allele count of 1, and a minor allele frequency of 0.01 (SNP present in at least three individuals). To ensure evenness across sequences, read depth was calculated via Plink v1.9 and SNPs with a depth between 10 and 40 reads per sequence (a minimum of half and a maximum of twice the average coverage of samples) were retained. Finally, Plink was used to remove all loci in linkage disequilibrium using a window size of 50 kb, step size of 10 bp and an r^2^ threshold of 0.1. The final dataset of 1,004,498 high quality SNPs was used for all population genomic analyses unless otherwise stated.

Genetic clusters (*K*) were identified with the program fastSTRUCTURE v1.0 (Raj et al., 2014) using a variational Baysian framework. We tested *K* = 1-10, with 10,000 iterations for each run and the priors set to be logistic. The “chooseK.py” script was implemented to determine the best estimate of *K* and a STRUCTURE-style plot generated using the base functions of R v4.2.3 (R Core Team, 2025).

Principal components analysis (PCA) was performed using Plink and plotted in R to visualise genetic differentiation between sites. Genetic differentiation was quantified as Wright’s *F*_ST_ calculated via Plink v2.00a5 (Chang et al., 2015) using a jackknife block size of 1,000 iterations to calculate the standard error around the estimate.

#### 2. Genomic diversity

Individual genomic diversity was calculated as autosomal observed heterozygosity (H_O_) as recommended by Schmidt et al. (2021). So as not to bias genotype calls to only biallelic loci and the two most common alleles within samples, individual autosomal H_O_ was calculated from each alignment file. In short, the number of heterozygous variants (compared to the reference genome) was divided by the size of the reference genome (3192.63 Mb). Individual autosomal H_O_ values were divided by the average autosomal H_O_ across individuals to calculate standardised heterozygosity (H_S_) (Coulon, 2010). H_S_ is a useful metric to place the genomic diversity of individuals into context with the samples included, as the average H_S_ is one and individuals with an H_S_ value above one are more diverse than average and individuals with an H_S_ below one are less diverse than average. Average H_S_ (and standard errors) was calculated per site and subsite.

#### 3. Summary statistics

To be consistent with calculations for individual SNP based H_O_, per-loci observed and expected (H_E_) heterozygosity was calculated with only a 100% call rate filter applied on the joint genotyping file using VCFtools. Nucleotide diversity (π) was calculated from the full quality filtered SNP set (*N* = 1,004,498) using windows of 10,000 in VCFtools. Average (with associated standard errors) per-loci H_O_, H_E_ and π were calculated for each site and subsite.

To estimate whether loci are under selection across the genome, we calculated Tajima’s D in 10,000 sliding windows via VCFtools. Tajima’s D was calculated separately for each site and subsite as this statistic assumes no population structure among samples. Average Tajima’s D (and associated standard error) was calculated for each site and subsite.

#### 4. Runs of homozygosity

From the unfiltered joint genotyping file, we removed all identified putatively sex-linked scaffolds to ensure that male heterosomes would not influence estimates of ROH. Next, we filtered on a call rate of 1 as missing data can greatly influence ROH estimates (Duntsch et al., 2021). The SNP dataset for ROH was not filtered for linkage disequilibrium or minor allele frequencies as per Meyermans et al. (2020). Following the removal of sex-linked scaffolds and missing data, the SNP dataset for ROH analysis included 15,187,654 SNPs.

We ran Plink v1.9 to detect ROH using a sliding window of 50 SNPs, requiring a minimum size of 100 kb and comprising at least 100 SNPs to be considered a ROH. One heterozygous SNP was allowed per window to account for genotyping errors. A minimum of one SNP per 50 kb was required to call a ROH to ensure that variable coverage across the genome did not bias detection. The maximum gap allowed between two SNPs to be considered adjacent ROH was set to 200 kb. Finally, at least 5% of windows were required to contain a homozygous SNP to be considered within a ROH. ROH were binned into size classes of > 100 kb, 500 kb, and 1 Mb.

ROH-based inbreeding coefficients (*F*_ROH_) were calculated based on the proportion of the genome present in ROH. As not all of the genome is considered in the ROH detection analysis (e.g., scaffolds shorter than the minimum ROH length are not considered), we calculated the maximum ROH our analysis could detect (i.e., the genome coverage for ROH) using a simulated individual with a completely homozygous genotype following Meyermans et al. (2020). We then calculated *F*_ROH_ as the sum of all ROH in an individual divided by the maximum ROH length our analysis could detect (3,080,060 kb; 96% of the reference genome). *F*_ROH_ was calculated for all ROH > 100 kb, 500 kb, and 1 Mb to enable comparisons of inbreeding coefficients across studies.

To investigate the association of ROH with the presence of *Chlamydia* shedding in koalas, we identified long ROH (> 1 Mb) that were only present in individuals that were positive for a *Chlamydia* infection compared to individuals with no detectable infection at the time of sampling across the *Chlamydia*-affected sites only, i.e. Richmond Range, Port Macquarie, Upper Nepean (within Woronora Plateau), and the Southern Tablelands. Of the long ROH unique to chlamydia-positive koalas, we identified the genes the ROH overlapped with (as annotated by FGENESH++) using bedtools intersect v2.27.1 (Quinlan & Hall, 2010). We retained genes that overlapped in at least 17 individuals (15% of individuals with a positive *Chlamydia* infection, roughly half the number of individuals sequenced at each site; (Silver et al., 2025) and ran a blastp search for these genes against the UniProt nonredundant protein database, retaining the top result for each protein. To determine protein function, we ran a Gene Ontology (GO) analysis using GONet (Pomaznoy et al., 2018) where the resulting GO terms were clustered and visualised using REViGO (Supek et al., 2011).

#### 5. Inbreeding and relatedness

We calculated individual inbreeding coefficients (*FH*) via Plink. *FH* is the probability that two alleles are identical by descent, originating from a common ancestor (Charlesworth, 2003). *FH* values were averaged and the associated standard error calculated for each site and subsite.

Relatedness between individuals was measured and presented as mean kinship (MK), which is the average kinship of one individual to all individuals included in the analysis (including self) and is relative to the samples included in the analysis. MK can be used to detect first- and second-order relationships, and show which sites are more related to themselves than others. We first calculated the triadic maximum likelihood estimator (TrioML) (Wang, 2007) on a random subset of 10,000 SNPs, due to computational requirements, in COANCESTRY v1.0 and divided this value by two to obtain MK. Average MK values, and associated standard errors, within sites and subsites and between sites and subsites were calculated.

Increased levels of inbreeding reduce genome-wide diversity and may potentially cause a loss of alleles responsible for disease resilience (Hedrick & Garcia-Dorado, 2016). The mean *FH* of koalas with a positive *Chlamydia* infection at the time of sampling was compared to the mean *FH* of those with a negative *Chlamydia* shedding result across *Chlamydia*-affected sites (see above) using an independent sample *t-test*. We used *FH* for this comparison as it is calculated on a per-locus basis (Chang et al., 2015), as opposed to *F*_ROH_ which is the proportion of the genome in ROH and may under-represent how much of the genome is identical by descent if these loci are not within a ROH.

#### 6. Gene flow

We calculated gene flow between sites and subsites, using a random subset of 50,000 SNPs from the filtered joint genotyping file, using the *DivMigrate* function in the *diveRsity* R package (Sundqvist et al., 2016). *DivMigrate* calculates genetic differentiation, based on allele frequencies, between populations to determine directional genetic divergence between pairs of sites. The estimates include uni- and bi-directional gene flow. Here genetic differentiation was measured as the effective number of migrants (*Nm*) (Alcala et al., 2014) with 1,000 bootstrap iterations to test for statistical significance in asymmetrical gene flow. We applied a minimum threshold of *Nm* > 0.3 to remove low rates of shared allele frequencies which are unlikely to represent meaningful genetic exchange.

#### 7. Effective population size

Contemporary effective population size (N_E_) was calculated using GoNe (Santiago et al., 2020). Putatively sex-linked scaffolds were removed from the fully filtered joint genotyping file resulting in 971,166 SNPs retained for analysis. As GoNe is heavily impacted by population structure/admixture (Novo et al., 2023), we ran GoNe for each sampling location as these were determined as individual genetic clusters by the *fast*STRUCTURE analysis (see Results). There is a known artefact in GoNe analyses, caused by population structuring, which results in a steep increase in N_E_ followed by a sudden drop. This artefact can be reduced when applying the most appropriate value for the maximum recombination rate parameters (abbreviated as hc). We therefore ran multiple iterations of GoNe to determine the most appropriate hc (0.01, 0.05 or 0.1) and across different recombination rates (cM/Mb). A previous recombination map built using over 400 koala sequenced genomes (Kovacs et al., 2025) showed that recombination rates ranged from 0 to 0.2 cM/Mb for NSW koalas. As such, we tested cM/Mb values of 0.01, 0.05, 0.1, 0.15, and 0.2. The optimal combination which reduced the potential artefacts from population structuring was 0.15 cM/Mb and 0.01 hc. To obtain confidence intervals around our estimate, we performed jack-knifing using the optimal cM/Mb and hc combination, where we iteratively removed one individual at a time and randomly subsampled SNPs in each replicate (a maximum of 50,000 SNPs per scaffold).

### Identification of candidate loci associated with Chlamydia infection

To identify potential candidate loci associated with a *Chlamydia* shedding in koalas, we performed genome-wide association studies (GWAS) using three methods (as per (Batley et al., 2021; Silver et al., 2022)), to reduce the chance of false positives in our GWAS. Slightly different filtering was applied for the subsequent GWAS compared to the population genomic statistics. The unfiltered joint genotyping file was filtered at a call rate of 1, using only biallelic sites, a minor allele count of 1, a minor allele frequency of 0.01, read depth between 10 and 40 via VCFtools, removing sites not in Hardy-Weinberg equilibrium (HWE; p < 1×10^-6^) and removing sites in linkage disequilibrium via Plink2 resulting in a final SNP set of 817,447 loci. Noting two versions of Plink were used because not all flags are available in both versions.

We performed GWAS using Plink with both a chi-square and Fisher’s exact test and retained only highly significant loci from these analyses (p < 0.001) (Batley et al., 2019; Batley et al., 2021; Silver et al., 2022; Wright et al., 2020). We supplemented these GWAS analyses with a Weir and Cockerham’s *F*_ST_ test (Weir & Cockerham, 1984) in VCFtools. Loci with a standard deviation of more than five from the mean were retained as significant (Axelsson et al., 2013; Batley et al., 2021; Silver et al., 2022). Any loci that were identified as significant in two or more of these tests were retained for further analysis.

As disease susceptibility is a complex trait, we implemented a multi-locus test on the significant loci identified in the GWAS (*N* = 4, 953). We performed a Random Forest analysis following Brieuc et al. (2018) where we first corrected for population structure and relatedness among samples. We performed two-proportion *z-tests* to detect differences in the mean ages and sex ratios between positive and negative *Chlamydia* shedding individuals. Neither age nor sex were significantly different between the case (positive *Chlamydia* shedding) and control (negative *Chlamydia* shedding) groups and so were not included as covariates in the Random Forest analysis (see Results). The number of case versus control individuals was uneven (*N* = 115 and 38 respectively). We corrected for this imbalance by balancing the representation of case and control individuals within the training data. To do this, we over-sampled the under-represented class (here controls) and under-sampled the over-represented class (here cases). We set the “sampsize” parameter of the “randomForest” function to be two-thirds of the total number of controls (*N* = 25) which randomly selects 25 individuals from the case and control groups for each iteration of the Random Forest run (Brieuc et al., 2018). We then iteratively tested the most appropriate values for *mtry* and *ntree* that resulted in the lowest out-of-bag error rate (OOB-ER) by running subsets of predictor loci (p) of √p, 2 x √p, 0.1p, 0.2p, p/3, and p and a range of *ntree* values from 100 to 200,000. To select loci predictive of *Chlamydia* infection from our Random Forest analysis, we employed a back-purging approach which consists of running multiple Random Forest runs on a slightly larger subset of the most important loci (a candidate group of loci that minimise OOB-ER) and recording the OOB-ER for each run. The locus with the highest OOB-ER is dropped iteratively until only two loci remain. The best set of loci for which OOB-ER is minimised is then identified and said to be predictive of chlamydia infection.

Our Random Forest analysis identified 34 loci associated with *Chlamydia* shedding (see Results). ANNOVAR was used to determine the genomic position of these loci (intergenic, intronic or exonic) associated with *Chlamydia* shedding. To determine protein function, we ran a GO analysis using GONet for all intronic loci where the resulting GO terms were clustered and visualised using REViGO. Allele frequency for the 34 candidate loci were calculated between case and control cohorts as well as between sites and subsites, including those where *Chlamydia* shedding was not detected in sampled koalas at the time of sampling, using VCFtools and visualised as a heatmap using ggplot2 (Villanueva & Chen, 2019).

## Results

For the 259 koalas included in this study, the average whole genome re-sequencing coverage was 22.23 (range: 13.45 – 32.61). Of these individuals, 115 tested positive for *Chlamydia* shedding at the time of sampling and 38 had no *Chlamydia* shedding detected (considered negative at the time of sampling). These individuals were from the sites currently impacted by *Chlamydia* (Richmond Range: *N* = 20 positive and *N* = 9 negative; Port Macquarie *N* = 41 positive and *N* = 15 negative; Woronora Plateau [Upper Nepean]: *N* = 8 positive and *N* = 8 negative; and Southern Tablelands: *N* = 46 positive and *N* = 6 negative) and 106 individuals were from sites where *Chlamydia* was not detected in sampled koalas (Narrandera: *N =41*; Kanangra Boyd: *N=40*, Woronora Plateau [Georges River]: *N=25*).

Per the *fastSTRUCTURE* analysis, the optimum value of *K* was six, corresponding to the six sampling sentinel sites with minimal overlap between sites as per the PCA (Figure 1b-c). Except for Narrandera, the greatest degree of variance between sites (PC1) appears to be explained by geographic distance from north to south (Figure 1b). The only sites to exhibit admixture were between Kanangra-Boyd, Woronora Plateau and the Southern Tablelands, which may likely reflect historical movement (Figure 1c), with these same sites clustering in the PCA with minimal overlap observed (Figure 1b). While koalas from Narrandera appear to represent a single cluster, there are some individuals that appear to have a small proportion of Richmond Range’s ancestry (Figure 1c). Previous work has shown that Narrandera animals were sourced from both Victoria and the area around south-east Queensland and northern NSW (McLennan et al., 2025a). Pairwise *F*_ST_ results were significant (95% confidence intervals did not encompass zero) across all pairs of sites and subsites, with the higher *F*_ST_ values between geographically distance sites (Table 1).

**Table 1.**
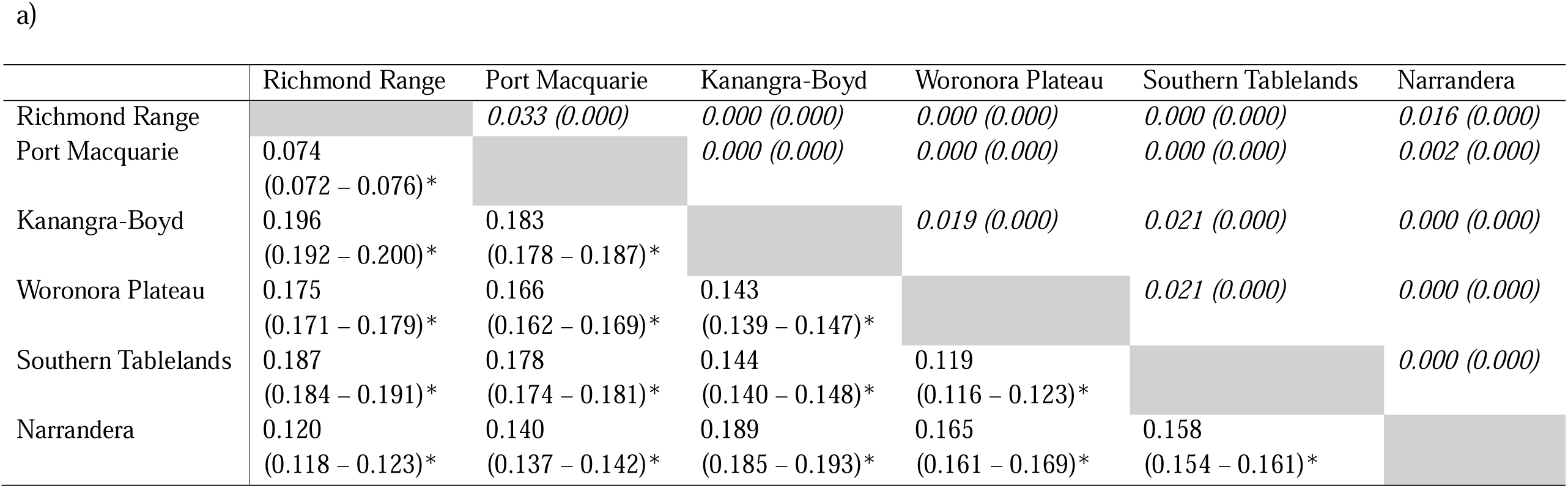
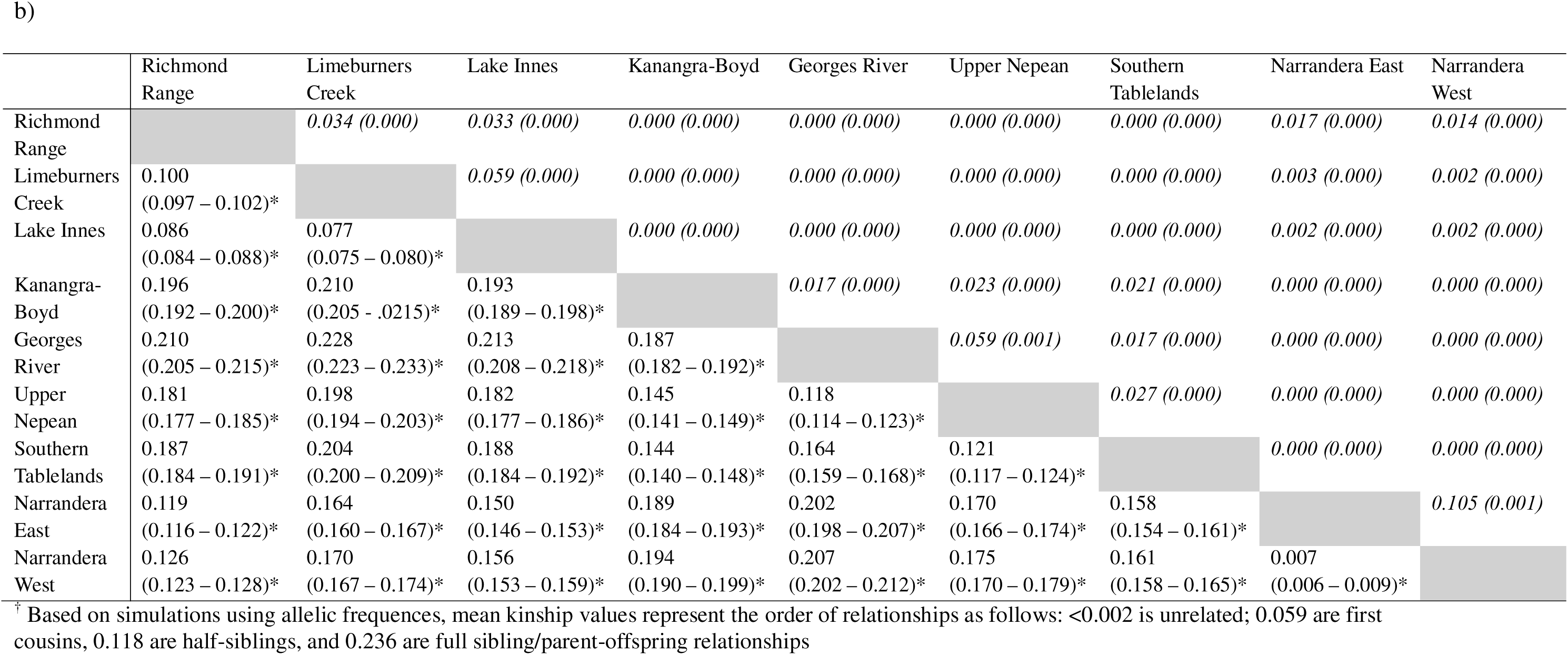
Pairwise population differentiation (*F_ST_*) and associated 95% confidence intervals between sampling sites (a) and sampling subsites (b) (below diagonal; grey; statistical significance was inferred at *p* < 0.05 and indicated with *); average mean kinship (MK^†^) with standard errors (SE) between sites (in italics, above diagonal)

H_S_ between sentinel sites varied, with Richmond Range and Port Macquarie representing the highest average H_S_ and Kanangra-Boyd and Woronora Plateau the lowest (Figure 1d), with the Georges River subsite likely driving the lower Woronora Plateau (Table 2). There were different levels of H_S_ between some subsites where Lake Innes koalas had higher H_S_ than Limeburners Creek (site: Port Macquarie) and similarly Upper Nepean koalas had higher H_S_ than those from Georges River (site: Woronora Plateau) (Table 2; Figure 1d). Koalas across the Narrandera subsites had similar levels of H_S_ (Table 2; Figure 1d). Patterns of H_O_ and H_E_ followed H_S_ values as did π (Table 2). Tajima’s D values were positive across all sites, indicative of either balancing selection or population bottlenecks (Table 2).

**Table 2.** Summary statistics for the 259 koalas included in this study including SNP-based observed heterozygosity (SNP H_O_) and expected heterozygosity (SNP H_E_), autosomal heterozygosity (Autosomal H_O_), standardised heterozygosity (H_S_), individual inbreeding (*FH*), proportion of the genome in runs of homozygosity longer than 1 mb (*F_ROH_*), mean kinship (MK), nucleotide diversity (π), Tajima’s D, and effective population size (N_E_)

| Site | SNP $H_O$<br>(± SE) | SNP $H_E$<br>(± SE) | Autosomal<br>$H_O$ (± SE) | Autosomal<br>$H_S$ (± SE) | $FH$ (± SE) | $F_{ROH}$ (±<br>SE) | MK<sup>†</sup> (± SE) | $\pi$ (± SE) | Tajima's D | $N_E$<br>(95% CI) |
| --- | --- | --- | --- | --- | --- | --- | --- | --- | --- | --- |
| Richmond Range | 0.255<br>(0.001) | 0.243<br>(0.001) | 2.31x10<sup>-3</sup><br>(8.99x10<sup>-6</sup>) | 1.173<br>(0.005) | -0.092<br>(0.003) | 0.003<br>(0.001) | 0.098<br>(0.002) | 8.24x10<sup>-5</sup><br>(2.47 x10<sup>-7</sup>) | 0.438<br>(0.004) | 214.477<br>(147.834–<br>281.120) |
| Port Macquarie | 0.229<br>(0.001) | 0.235<br>(0.001) | 1.88x10<sup>-3</sup><br>(9.50x10<sup>-6</sup>) | 1.030<br>(0.005) | -0.002<br>(0.004) | 0.007<br>(0.001) | 0.088<br>(0.001) | 7.90x10<sup>-5</sup><br>(2.48 x10<sup>-7</sup>) | 0.576<br>(0.004) | 340.093<br>(312.041–<br>368.145) |
| *Limeburners Creek* | 0.223<br>(0.001) | 0.211<br>(0.001) | 1.88x10<sup>-3</sup><br>(1.38x10<sup>-5</sup>) | 1.003<br>(0.008) | 0.011<br>(0.005) | 0.008<br>(0.001) | 0.135<br>(0.002) | 7.98x10<sup>-5</sup><br>(2.64 x10<sup>-7</sup>) | 0.567<br>(0.004) |  |
| *Lake Innes* | 0.233<br>(0.001) | 0.225<br>(0.001) | 1.91x10<sup>-3</sup><br>(9.91x10<sup>-6</sup>) | 1.046<br>(0.005) | -0.012<br>(0.004) | 0.007<br>(0.001) | 0.108<br>(0.001) | 7.89x10<sup>-5</sup><br>(2.54 x10<sup>-7</sup>) | 0.518<br>(0.004) |  |
| Kanangra-Boyd | 0.203<br>(0.001) | 0.195<br>(0.001) | 1.66x10<sup>-3</sup><br>(2.44x10<sup>-5</sup>) | 0.908<br>(0.013) | 0.128<br>(0.011) | 0.038<br>(0.003) | 0.163<br>(0.001) | 7.11x10<sup>-5</sup><br>(2.47 x10<sup>-7</sup>) | 0.586<br>(0.004) | 313.648<br>(258.472–<br>368.824) |
| Woronora Plateau | 0.204<br>(0.001) | 0.213<br>(0.001) | 1.63x10<sup>-3</sup><br>(2.04x10<sup>-5</sup>) | 0.896<br>(0.011) | 0.088<br>(0.009) | 0.025<br>(0.001) | 0.118<br>(0.002) | 7.55x10<sup>-5</sup><br>(2.48 x10<sup>-7</sup>) | 0.609<br>(0.004) | 80.988<br>(60.071–<br>94.905) |
| *Georges River* | 0.194<br>(0.001) | 0.182<br>(0.001) | 1.56x10<sup>-3</sup><br>(2.14x10<sup>-5</sup>) | 0.855<br>(0.012) | 0.121<br>(0.008) | 0.031<br>(0.001) | 0.187<br>(0.002) | 7.22x10<sup>-5</sup><br>(2.61 x10<sup>-7</sup>) | 0.439<br>(0.003) |  |
| *Upper Nepean* | 0.220<br>(0.001) | 0.206<br>(0.001) | 1.74x10<sup>-3</sup><br>(1.48x10<sup>-5</sup>) | 0.960<br>(0.008) | 0.037<br>(0.009) | 0.017<br>(0.001) | 0.138<br>(0.002) | 7.92x10<sup>-5</sup><br>(2.62 x10<sup>-7</sup>) | 0.530<br>(0.003) |  |
| Southern Tablelands | 0.215<br>(0.001) | 0.209<br>(0.001) | 1.80x10<sup>-3</sup><br>(1.09x10<sup>-5</sup>) | 0.986<br>(0.006) | 0.096<br>(0.004) | 0.018<br>(0.001) | 0.127<br>(0.001) | 7.29x10<sup>-5</sup><br>(2.41 x10<sup>-7</sup>) | 0.687<br>(0.004) | 4262.280<br>(3535.250–<br>4989.310) |
| Narrandera | 0.241<br>(0.001) | 0.233<br>(0.001) | 1.91x10<sup>-3</sup><br>(1.55x10<sup>-5</sup>) | 1.048<br>(0.008) | -0.005<br>(0.006) | 0.018<br>(0.001) | 0.109<br>(0.001) | 7.90x10<sup>-5</sup><br>(2.48 x10<sup>-7</sup>) | 0.666<br>(0.004) | 2290.800<br>(0–<br>37456.371) |
| *East* | 0.242<br>(0.001) | 0.231<br>(0.001) | 1.92x10<sup>-3</sup><br>(1.86x10<sup>-5</sup>) | 1.054<br>(0.010) | -0.010<br>(0.007) | 0.018<br>(0.001) | 0.109<br>(0.002) | 8.06x10<sup>-5</sup><br>(2.54 x10<sup>-7</sup>) | 0.585<br>(0.004) |  |
| *West* | 0.239<br>(0.001) | 0.225<br>(0.001) | 1.90x10<sup>-3</sup><br>(2.63x10<sup>-5</sup>) | 1.041<br>(0.014) | 0.003<br>(0.009) | 0.018<br>(0.001) | 0.118<br>(0.002) | 8.04x10<sup>-5</sup><br>(2.56 x10<sup>-7</sup>) | 0.528<br>(0.003) |  |

<sup>†</sup> Based on simulations using allelic frequencies, mean kinship values represent the order of relationships as follows: <0.002 is unrelated; 0.059 are first cousins, 0.118 are half-siblings, and 0.236 are full sibling/parent-offspring relationships

The number of ROH increased from north to south, with Richmond Range (average number of ROH = 2505.76) having the fewest runs and Woronora Plateau (average number of ROH = 4194.32) the most (Table S1). *F*_ROH_ also decreased from north to south, however Kanangra-Boyd had the highest *F*_ROH_ for runs longer than 1 Mb (Figure 2a). Similarly, ROH longer than 1 Mb increased from Richmond Range through to Narrandera with Kanangra-Boyd, Woronora Plateau, the Southern Tablelands, and Narrandera showing consistent ROH locations across scaffolds (Figure S1). When comparing ROH longer than 1 Mb between *Chlamydia* positive and *Chlamydia* negative koalas, there were four ROH that were present in at least 15% (*N* = 17) of *Chlamydia* positive koalas and not in *Chlamydia* negative individuals (Figure 2b). These four ROH were located along Scaffold two and four and intersected with 18 genes. Of these 18 genes, 12 had annotations in the UniProt database (Table S2). Four of these genes have known associations and/or differential expression profiles in humans infected with *C. trachomatis* (Table S2). The 12 annotated genes were enriched in several GO pathways including chromosome segregation, immune system and reproductive processes, cell adhesion, cell division, and response to stress (Table S3; Figure S2).

**Figure 2.**
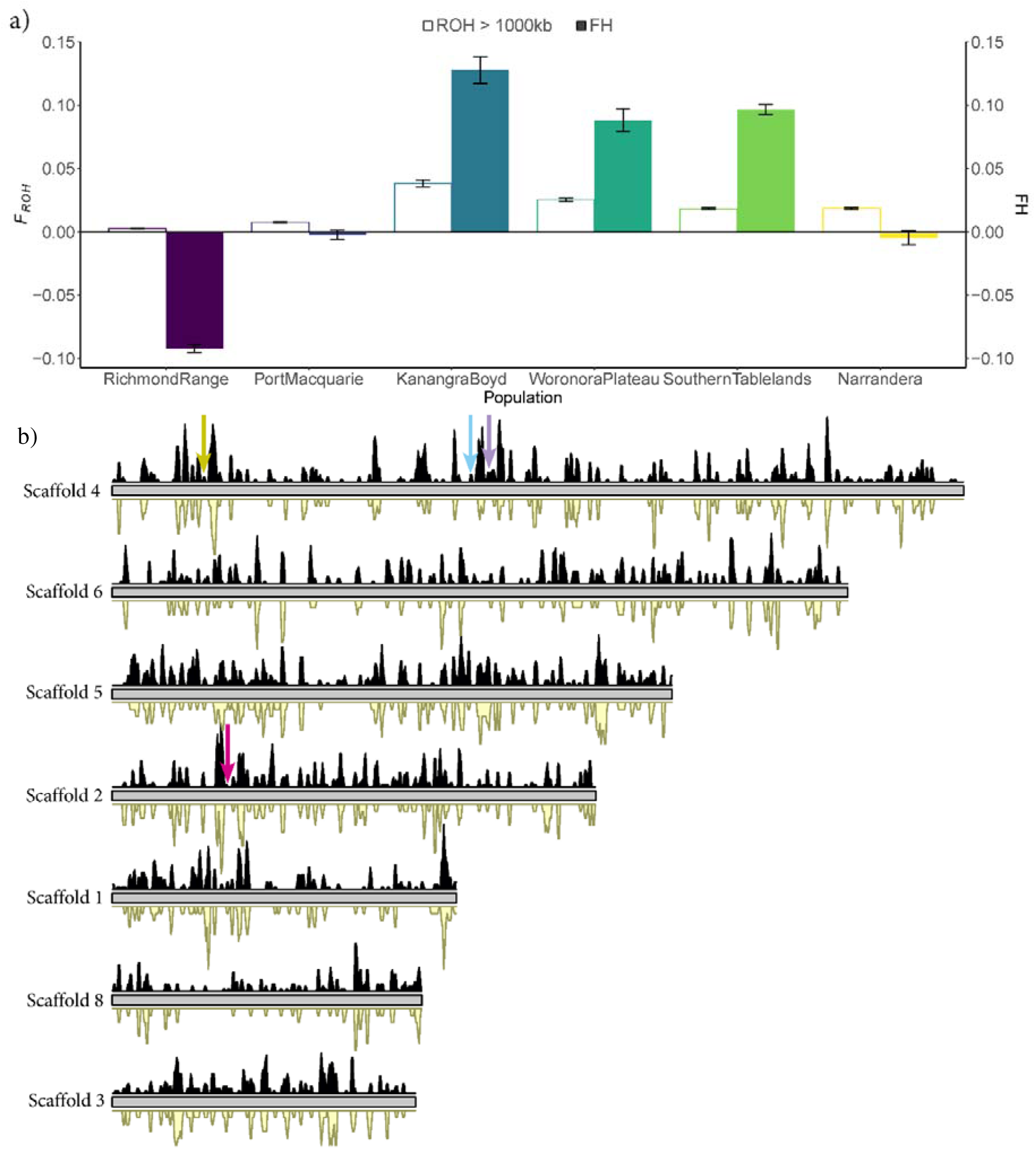
a) Average proportion of the genome in runs of homozygosity (*F*_ROH_) longer than 1,000 kb (outline bars) and average individual inbreeding (*FH;* coloured bars) across the New South Wales Sentinel Program sites, error bars represent standard error; b) runs of homozygosity (ROH) longer than 1,000 kb in individuals that tested positive for a chlamydia infection at the time of sampling (black) and those that tested negative (yellow) across the seven longest scaffolds representing the autosomes of the koala genome, ordered from longest to shorted scaffold. The height of the peak shows the density of the ROH across the scaffolds, higher peaks represent more ROH in a region. The arrows represent ROH present only in chlamydia positive individuals (*N* > 17 koalas) encompassing the following genes of un-annotated (yellow arrow), COL4A3 (blue arrow), ZSWIM6, SMIM15, NDUFAF2, ERCC8, RPL14, ELVL7, DEPDC1B (purple arrow), and EpCAM, MSH2, KCNK12, MSH6 (pink arrow). Gene Ontology pathways for these genes presented in Table S3.

Average *FH* was highest for Kanangra-Boyd, reflective of this population having the highest *F*_ROH_ longer than 1 Mb, followed by the Southern Tablelands and Woronora Plateau (Table 2; Figure 2b). Similar to the H_S_ results, *FH* was different between the Port Macquarie and Woronora Plateau subsites with Limeburners Creek having higher *FH* than Lake Innes and Georges River higher than Upper Nepean (Table 2). Average *FH* was slightly higher in Narrandera West compared to Narrandera East (Table 2). Mean *FH* was significantly higher in *Chlamydia* positive koalas (average *FH* = 0.026) at the time of sampling compared to *Chlamydia* negative individuals (average *FH* = −0.0051) (*t* = −2.31, df = 151, p-value = 0.022). Average MK of all individuals within the sample set (all six sentinel sites) was higher than 0.118 (half-sibling relationships) within Kanangra-Boyd, the Southern Tablelands and the Woronora Plateau, including subsites of Georges River and Upper Nepean, and higher than first cousins (MK > 0.059) within Narrandera and Woronora Plateau (Table 2). There was no detected MK between most sites, however there was low MK (< 0.059; first-cousins) between Richmond Range and Port Macquarie, between Woronora Plateau and Kanangra-Boyd, and between the Southern Tablelands, Kanangra-Boyd and Woronora Plateau (Table 1a). Similarly, there was low MK (< 0.059) between most subsites, except for between Narrandera East and West which was approaching half-sibling relationships on average (Table 1b). The average MK within each site (i.e. only individuals within the site were included in the analysis) was lower than when all sites were included, with Woronora Plateau exhibiting a cousin relationship (Table S4).

Contemporary *N*_E_ was high for the Southern Tablelands and Narrandera, moderate for Richmond Range, Port Macquarie and Kanangra-Boyd and notably low for Woronora Plateau (Table 2). Gene flow was minimal between sites and subsites (Figure 3). High gene flow (*Nm* > 0.7) was only present between the subsites of Narrandera East and West, which are geographically closer than all other subsites (Figure 3b). There was no medium gene flow (< 0.7 *Nm* > 0.5) between any site or subsite and low gene flow (< 0.5 *Nm* > 0.3) only between the subsites of Limeburners Creek and Lake Innes (Port Macquarie) (Figure 3).

**Figure 3.**
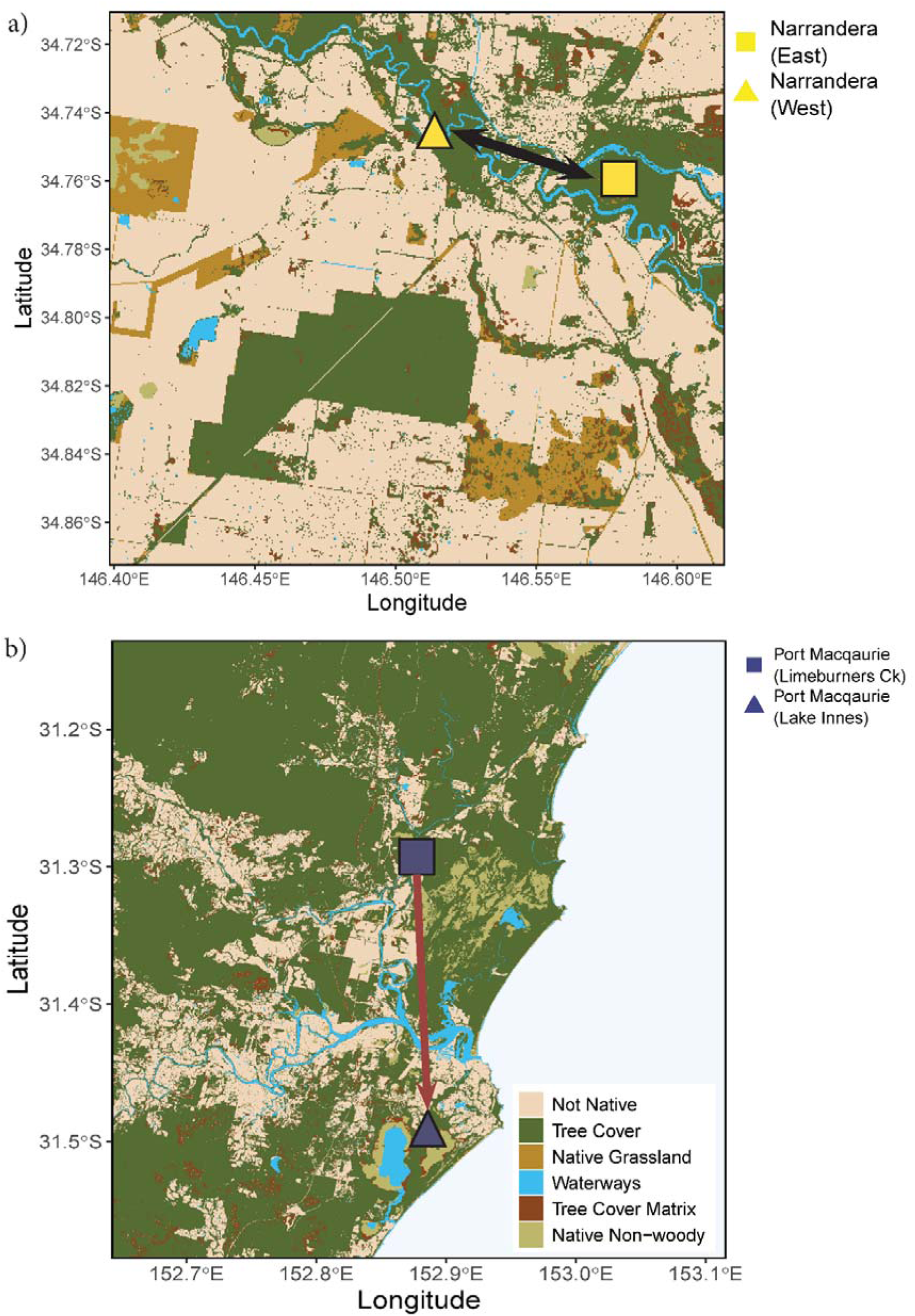
a) High gene flow (Number of effective migrants [*Nm*] > 0.7) between the subsites of Narrandera East and West plotted over available vegetation where Tree Cover and Tree Cover Matrix are considered suitable koala habitat, and b) low gene flow (*Nm* <0.5 and >0.3) between the subsites of Limeburners Creek and Lake Innes in Port Macquarie.

Neither age (χ^2^ = 1.62, df = 1, p-value = 0.20) nor sex (χ^2^ = 0.067, df = 1, p-value = 0.80) were significantly different between the case and control groups in the GWAS analysis. The optimal combination of *mtry* and *ntree* was √p (70 loci from full 4,953 input set) and 200,000 respectively to achieve the lowest OOB-ER and repeatability (correlation of >95% of the importance values between replicate Random Forest runs). The most important candidate group of loci that minimised OOB-ER was the top 5%, therefore the back-purging approach was performed on the top 10% of candidate loci. The back-purging approach resulted in 34 loci identified as candidates for predicting *Chlamydia* shedding in koalas. Of these 34 loci, ANNOVAR analysis identified 15 of these in intronic regions, three loci upstream of a gene, and the remaining 16 loci in intergenic regions, ranging from 1kb to 739kb away from a gene (Table S5). No candidate loci were within exons. Of the intronic loci, nine were associated with annotated genes in the Uniport database (Table S5). Of the loci upstream of genes, only one gene was annotated (Table S5). Of the loci in the intergenic regions, 11 had at least one gene annotated (Table S5). The intronic and upstream loci associated with annotated genes were enriched for several GO pathways including the same as those identified for the genes in ROH in *Chlamydia* infected individuals such as cell motility, reproductive and immune system processes, chromosome segregation, cell division, cell adhesion, and response to stress (Table S6; Figure S3). The allele frequencies of the 34 candidate loci that were detected from the GWAS of *Chlamydia* positive and *Chlamydia* negative individuals varied across populations (Figure 4). Consistently, the sites and subsites within Port Macquarie (*Chlamydia* present site), Kanangra-Boyd (*Chlamydia* absent site), Woronora Plateau (Upper Nepean, (*Chlamydia* present site; Georges River *Chlamydia* absent site) and the Southern Tablelands (*Chlamydia* present site) had similar allele frequencies at these 34 candidate loci to the *Chlamydia* positive infected individuals from the GWAS analyses (Figure 4). Conversely, Richmond Range (*Chlamydia* present site) had similar allele frequencies to the *Chlamydia* negative individuals, whilst Narrandera subsites (*Chlamydia* absent site) had alleles that were consistent with both the *Chlamydia* positive and negative individuals (Figure 4).

**Figure 4.**
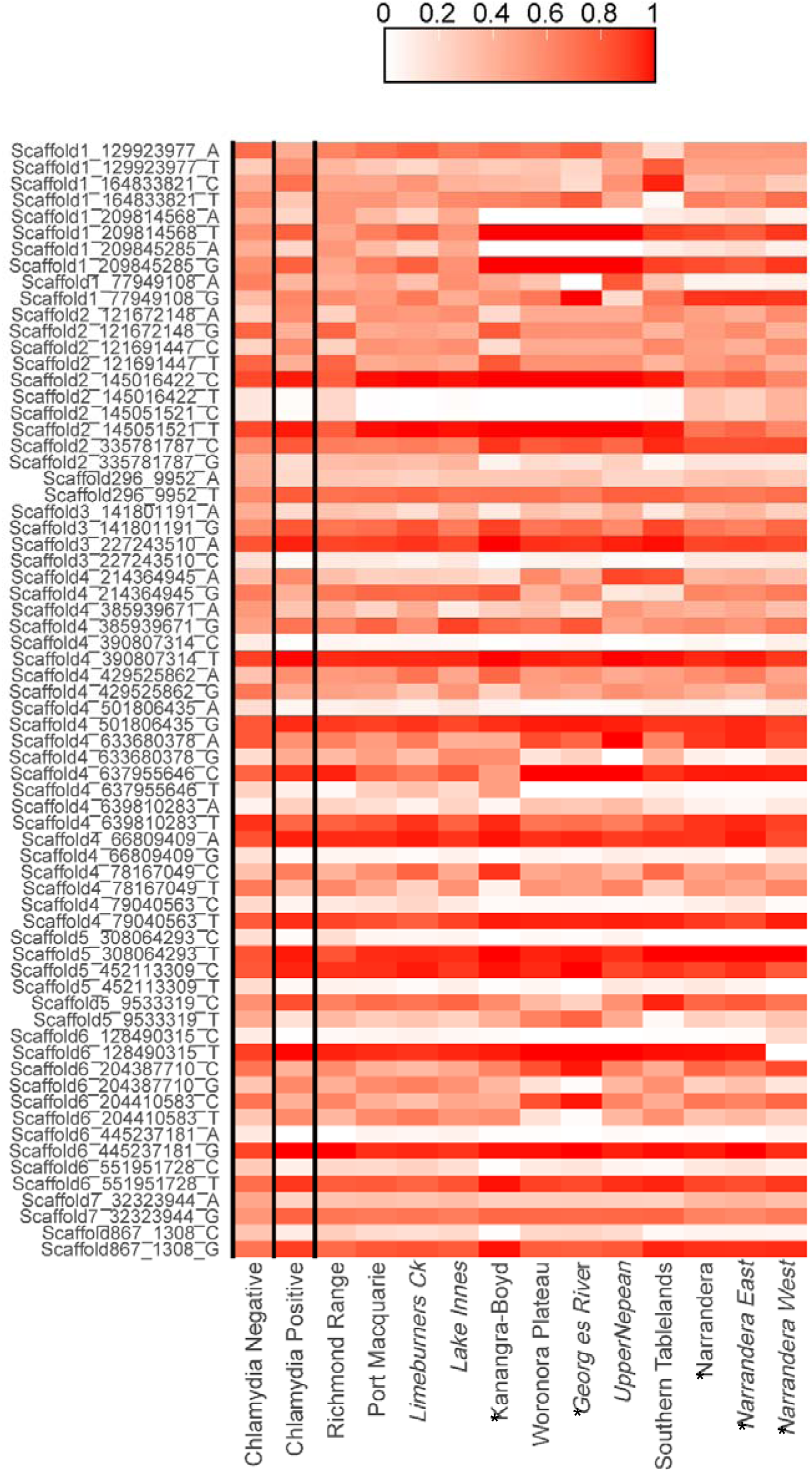
Heatmap of allele frequencies for the 34 candidate loci identified with a Random Forest analysis in the *Chlamydia* positive (N=115) and *Chlamydia* negative (N=38) individuals. Allele frequencies for these same 34 candidate loci are also shown for each site (subsites in italics), highlighting that these alleles exist in populations that are known to be *Chlamydia* present and *Chlamydia* absent sites (*). Genomic position and allele provided in loci name (y-axis).

## Discussion

Our assessment of the population genomic health (diversity, inbreeding, relatedness, Ne) of the six sentinel populations showed a general trend of higher diversity and lower inbreeding in the northern sites compared to those southern sites, which aligns with previous work assessing the genomic status of koalas across eastern Australia (McLennan et al., 2025a). Although the gene flow analysis shows no contemporary gene flow between the six sentinel sites, the *fastStructure* analysis showed some admixture likely reflecting the historical movement of koalas as noted in McLennan et al. (2025a). Additionally, our finding of no contemporary gene flow is not surprising as the sites used in the program are geographically isolated. The increase in the number and length of ROH from the northern sites to the southern was expected and reflective of broader koala genomic patterns across eastern Australia (Johnson et al., 2018; McLennan et al., 2025a).

Here we show an association between individual levels of inbreeding and detection of *Chlamydia* shedding in koalas. That is, *Chlamydia*-positive individuals at the time of sampling had significantly higher inbreeding coefficients than *Chlamydia*-negative individuals regardless of their geographic location. We identified several genes in ROH that were unique to *Chlamydia* shedding koalas. Noting, this study has not examined the severity of disease resulting from *Chlamydia* infections. Mechanistic studies should follow up some of these genes to determine whether they may play a role in susceptibility to or severity of manifestation of chlamydiosis. Interestingly, similar degrees of inbreeding can be found in koala populations that may not yet have been challenged by this pathogen, with similar ROH profiles and candidate loci allele frequencies to *Chlamydia*-positive individuals. Our gene flow analysis supports the hypothesis that isolation rather than the presence of resilient genotypes and/or higher genomic diversity protects koala populations from chlamydiosis (McLennan et al., 2025a). While increasing avenues for gene flow to reverse inbreeding and subsequent genome-wide homozygosity may enable better resilience of populations to respond to *Chlamydia* infections, such actions may also provide pathways for chlamydial incursion into vulnerable populations (McLennan et al., 2025b). A thorough understanding of how genotypes and gene expression influence susceptibility to chlamydiosis is required to ensure the most appropriate management actions are taken. Future research should also investigate the environmental thresholds and landscape features that determine *Chlamydia* establishment and persistence, including stress physiology, habitat quality metrics, population density effects, and the geographic barriers that successfully protect isolated populations, such as Kanangra-Boyd and Narrandera, from pathogen introduction.

It is important to note that our comparisons, of both ROH and candidate loci, between *Chlamydia*-positive and *Chlamydia*-negative koalas occurred only at a single time-point. Even if individuals were *Chlamydia*-negative at the time of sampling, this does not necessarily mean they will remain so. Future work from the sentinel sites that utilises temporal data may better inform this. Similarly, variability of severity and clinical manifestation of disease expression remains a fertile source of further longitudinal studies. It is possible that some individuals may clear the infection with minimal long-term complications while others will sustain urogenital and ocular tissue damage leading to urinary complications, infertility and/or blindness (Quigley & Timms, 2020). The balance in host-pathogen outcomes for chlamydiosis for NSW koalas will be a key outcome of the longitudinal NSW Koala Sentinel Program.

While factors impacting inbreeding may also independently impact chlamydial transmission patterns or susceptibility, the relevance of the genetic loci identified in this study to mechanisms of chlamydial infection and persistence supports further investigation of their role as predisposing traits. Both the ROH and candidate loci analyses were enriched for several GO pathways such as cell motility, reproductive and immune system processes, chromosome segregation, cell division, cell adhesion, and response to stress. One ROH unique to *Chlamydia*-positive individuals encompassed the gene coding for the epithelial cell adhesion molecule EpCAM (Trzpis et al., 2007), and is likely to encompass its promoter region, with the ROH beginning more than 1,500 kb prior to the gene sequence (Clancy, 2008). Although induction of EpCAM by chlamydial infection likely influences development of disease by lessening the tight cell-cell adhesions and modulating cell proliferation and differentiation (Trzpis et al., 2007), at least in the ovarian bursa of the koala (Pagliarani et al., 2022) and fallopian tube of women (Kessler et al., 2012), its role during establishment and persistence of chlamydial infection is less studied. However, the role of EpCAM and associated modulation of NF-KB is studied in the context of neoplasia (Gaiser et al., 2012; Sankpal et al., 2013), demonstrating outcomes expected to be advantageous to chlamydial persistence: reduction of apoptosis, thus maintaining the infected cell as a host; and altered activation, migration and cytokine profile of epithelial cells; and antigen-presenting Langerhans cells, thus influencing the direction of subsequent local and systemic immune pathways. The expression of these genes, host regulated or *Chlamydia*-induced, would be well worth studying in the context of chlamydial infection in koalas.

Host cell DNA damage induced by *Chlamydia* spp. facilitates continued bacterial proliferation (Chumduri et al., 2013) by interfering with many defensive cell functions, including apoptosis. *C. trachomatis* in humans impedes DNA mismatch repair mechanisms by downregulating expression of MSH2, MSH6 and ERCC8 genes, which in healthy cells identify errors in the DNA sequence and corrects them during cell replication (Koster et al., 2022). Here we found ROH unique to *Chlamydia*-positive koalas encompassing the MSH2, MSH6, and ERCC8 genes and likely their promoter regions as well. As with the expression of EpCAM, further investigation is ongoing to determine whether homozygosity across these DNA repair genes is resulting in uniform, low expression, thus exacerbating DNA damage and favouring chlamydial persistence within the host cell.

Potential associations between immune genotypes/expression and *Chlamydia* presentation have been provided in the literature (Lau et al., 2014; Pagliarani et al., 2024; Quigley et al., 2018; Silver et al., 2022), but we only found one candidate locus loosely associated with an immune gene. This locus was situated in the intergenic region, so its regulatory influence is likely to be tenuous, between the ZFAT (669Kb) and CFRAP251 (252Kb) genes. The fact that we did not find any clear associations between, *Chlamydia* shedding and immune gene diversity may be due to the strong selective pressure placed upon immune genes to maintain diversity (Sommer, 2005). Indeed, in highly inbred species that have undergone severe bottlenecks, immune gene diversity is still maintained (Nelson et al., 2025). This lack of association between immune genes and *Chlamydia* shedding may also be attributed to the possibility that our *Chlamydia*-negative individuals simply had not yet been exposed to *C. pecorum*; or our use of a broad-scale genome-wide approach. It is possible that changes to inflammation and DNA repair mechanisms are more influential or predictive of *Chlamydia* infections than immune gene diversity. This opens several interesting avenues for future research beyond immune genes that may predispose koalas to progression of chlamydiosis through dysregulated inflammatory responses and/or DNA repair mechanisms and highlights the importance of generating whole genome data opposed to targeting specific genes.

Our findings reveal distinct genetic profiles associated with chlamydial infection across NSW koala populations, which will necessitate a tailored rather than a uniform conservation strategy. The interaction between genetic vulnerability and pathogen exposure creates distinct yet significant risk scenarios. Northern populations (Richmond Range and Port Macquarie) present the most significant management challenge, where high individual genetic diversity fails to prevent *Chlamydia* infections. This pattern suggests that challenge by *Chlamydia* leads to population-wide impact regardless of genetic diversity status, although individuals inbreeding coefficients may modulate individual outcomes within exposed populations. Further, the contrast between the Southern Tablelands and Kanangra-Boyd illustrates vulnerability and pathogen exposure. While inbreeding is similarly high, isolation from disease incursion may be the difference between low and high shedding rates. Longitudinal data will inform whether this is related to population viability. We therefore envisage that effective management will require a raft of population-specific interventions, including potentially intensive genetic rescue, *Chlamydia* or environmental management strategies for northern populations, and potentially strict biosecurity measures for current *Chlamydia*-free populations.

Disease resilience is multi-faceted with many cellular processes involved. Here we showcase the importance of employing a genome-wide approach to uncover hitherto unexplored mechanisms associated with chlamydial infection and persistence. Anthropogenic pressures leading to population isolation and subsequent inbreeding and loss of genomic diversity are hindering species’ capacity to co-adapt in the evolutionary arms race between hosts and pathogens. As such, overall conservation efforts should continue to focus on reduction of inbreeding and maintenance of diversity, however this study suggests that conservation measures that promote population connectivity may need to carefully consider the impacts of disease incursion. Increased understanding of the drivers of disease susceptibility remains essential in informing connectivity and translocation conservation measures for any threatened species, potentially helping to predict population disease vulnerability profiles and therefore levels of management intervention. This can only be achieved through investment into monitoring programs that permit longitudinal temporal studies. With species’ genetic diversity declining globally and wildlife diseases increasing, it is imperative to understand the underlying mechanisms of potential disease resilience and tailor conservation approaches to give populations a fighting chance at successful co-evolution.

## Acknowledgements

The authors acknowledge the traditional custodians of the land upon which koalas reside and pay our respect to their elders past and present. We thank the field team from the Taronga Conservation Society Australia for sample and metadata collection and curation, particularly Danielle Fryday and Elizabeth McConnell. Thank you as well to Kim Heasman, Konstanze Gebauer, Lauren Alexander, Hannah Newton and Rachel Jelic for their assistance with DNA extractions and sample management. We thank Swetansu Pattnaik from Illumina for bioinformatic support and the Ramaciotti Centre for Genomics for sequencing support. We thank Amazon Web Services and RONIN for providing compute resources. This work was conducted under the New South Wales Koala Sentinel Program funded by the New South Wales Department of Climate Change, Energy, Environment, and Water.

## Data availability

All raw sequence data for this project is publicly available through the New South Wales Government SEED, Central Resource for Sharing and Enabling Environmental Data in NSW (https://datasets.seed.nsw.gov.au/dataset/003600a1-d2d7-4761-8573-1a95212be7d0) and available to download from Amazon Web Services Open Datasets Program (https://koalagenomes.s3.ap-southeast-2.amazonaws.com/index.html#NSW_Sentinel/). Koala metadata is available on Dryad (private for peer review) XXX.

## Funding statement

This work was conducted under the New South Wales Koala Sentinel Program funded by the New South Wales Department of Climate Change, Energy, Environment, and Water.

## Conflict of interest statement

E.A.M. is a member of the National Koala Recovery Team Community Advisory Committee, an advisory committee to the Australian Federal Government. C.J.H. is a member of the New South Wales (NSW) Expert Panel for koalas, an advisory panel to the NSW Government, and K.B. was a member of this panel from 2015 to 2023. The authors declare they have no conflict of interest.

## Ethics approval statement

All koala capture and sampling was conducted under Animal Ethics Committee number 230718/02.

## Benefit-Sharing

All data produced as part of this project is publicly available, and findings of the research shared with the New South Wales Government Koala Monitoring and Research Team to inform conservation efforts.

## Author contributions

E.A.M., C.J.H., T.S.J.: conceptualisation. L.V.: sample and metadata collection. C.J.H., K.B., D.P.H., M. B.K., T.S.J.: resources. Z.C.: genome alignment and joint genotyping support. M.K.: chlamydia detection. A.C., B.R.W.: chlamydial detection. M.B.K., D.P.H.: chlamydial interpretation. E.A.M.: population genomic and genotype-disease association analyses with assistance from L.S. E.A.M.: data visualisation. E.A.M.: manuscript original draft. All authors edited the manuscript and reviewed the final submission for scientific content.

## Notes

https://koalagenomes.s3.ap-southeast-2.amazonaws.com/index.html#NSW_Sentinel/

