## Supplementary material for "Inbreeding and resultant homozygosity across key inflammation and DNA repair genes linked to chlamydial infection in New South Wales koalas"

**Table of Contents:**

| **Supplementary Methods** | Page 2 |
| --- | --- |
| **Table S1** | Page 3 |
| **Table S2** | Page 3 |
| **Table S3** | Page 4 |
| **Table S4** | Page 6 |
| **Table S5** | Page 6 |
| **Table S6** | Page 7 |
| **Figure S1** | Page 8 |
| **Figure S2** | Page 9 |
| **Figure S3** | Page 10 |

**Supplementary Methods**

If required, anaesthesia was supplemented with alfaxalone (0.8-1mg/kg) intravenously or intramuscularly or with isoflurane in oxygen administered via face mask. At the end of the procedure the medetomidine was reversed with atipamezole at 5 times the medetomidine dose, administered intramuscularly. Koalas were allowed a full hour of recovery post-anaesthesia prior to release. Morphometric and health measurements of weight, presence of pouch/back young, crown to tail length, head length, ear width and length, manus and pes length, age, body condition score, hydration, signs of cardiovascular, respiratory, dental, abdominal, musculoskeletal, urogenital, ocular, dermal, and lymph node abnormalities, evidence of ectoparasites, and chlamydia wet bottom score were collected for each individual. Tissue samples were taken from the ear pinna margin with a 4mm leather punch which was disinfected with F10 antiseptic (F10 Products Ltd, United Kingdom) and 70% ethanol prior to use. Fur around the punch site was clipped and the skin swabbed with a methanol/chlorhexidine soaked cotton ball then rinsed with 70% ethanol prior to the biopsy being taken. Pressure was applied to the punch site post-biopsy to staunch any bleeding. The tissue plug was then placed into 70% ethanol for storage at -80°C. For ocular samples, the swab was inserted into the dorsal conjunctival sac and gently rotated while moving it back and forth 5-10 times against the conjunctival mucosa, avoiding contact with the cornea. A separate swab was used for each eye. For the female urogenital sample, the cloaca is everted, and the swab gently inserted just above the clitoris and directed cranially into the urogenital sinus and rotated 5-10 times. For the male urogenital sample, the penis is extruded from the cloaca, extended, and the swab inserted into the urethra between the bifurcation of glans and rotated 5-10 times.

**Table S1** Runs of homozygosity (ROH) statistics for 259 wild New South Wales koala samples with sampling site and where relevant subsite provided. ROH are separated into sizes classes as well as the proportion of the genome in ROH (*F*_ROH_). Attached as an excel spreadsheet.

**Table S2** Details of the runs of homozygosity (ROH) that intersected with genes and were present in at least 17 chlamydia positive koalas and not in any chlamydia negative individuals including the scaffold (ROH scaffold), start (ROH start) and end (ROH end) position of the ROH, the scaffold in which the ROH intersected (Intersect scaffold), the database searched against (Database), the type of sequence (here all genes; Type), the gene start (Gene start) and end (Gene end) position, the strand orientation (Strand), the gene code in the annotation file (Gene code), the gene name identified via a blastp search (Gene name), whether the ROH was assumed to cover the promoter region of the gene (Promoter in ROH), the number of base pairs the ROH began prior to the gene sequence (BPs ROH starts before gene), whether links between genes and human’s infected with *Chlamydia trachomatis* have been made (Links to *Chlamydia trachomatis*) and links to the publication. Attached as an excel spreadsheet.

**Table S3** Gene Ontology (GO) terms and associated genes from 18 genes in runs of homozygosity between chlamydia positive and chlamydia negative koalas

| **GO term ID** | **GO term definition** | **No. of Genes** | **Genes** |
| --- | --- | --- | --- |
| GO:0007165 | Signal transduction | 5 | *COL4A3\|DEPDC1B\|EPCAM\|MSH2\|MSH6* |
| GO:0048856 | Anatomical structure development | 5 | *COL4A3\|EPCAM\|MSH2\|MSH6\|ZSWIM6* |
| GO:0034641 | Cellular nitrogen compound metabolic process | 5 | *ELOVL7\|ERCC8\|MSH2\|MSH6\|P50914* |
| GO:0008219 | Cell death | 3 | *COL4A3\|MSH2\|MSH6* |
| GO:0002376 | Immune system process | 3 | *EPCAM\|MSH2\|MSH6* |
| GO:0051276 | Chromosome organization | 3 | *ERCC8\|MSH2\|MSH6* |
| GO:0006259 | DNA metabolic process | 3 | *ERCC8\|MSH2\|MSH6* |
| GO:0030154 | Cell differentiation | 3 | *EPCAM\|MSH2\|ZSWIM6* |
| GO:0006950 | Response to stress | 3 | *ERCC8\|MSH2\|MSH6* |
| GO:0006810 | Transport | 2 | *KCNK12\|P50914* |
| GO:0007568 | Aging | 2 | *MSH2\|MSH6* |
| GO:0009056 | Catabolic process | 2 | *ERCC8\|P50914* |
| GO:0007155 | Cell adhesion | 2 | *COL4A3\|EPCAM* |
| GO:0009058 | Biosynthetic process | 2 | *ELOVL7\|P50914* |
| GO:0007049 | Cell cycle | 2 | *MSH2\|MSH6* |
| GO:0000003 | Reproduction | 2 | *MSH2\|MSH6* |
| GO:0040011 | Locomotion | 2 | *DEPDC1B\|EPCAM* |
| GO:0044403 | Symbiotic process | 2 | *MSH6\|P50914* |
| GO:0048870 | Cell motility | 2 | *DEPDC1B\|EPCAM* |
| GO:0006091 | Generation of precursor metabolites and energy | 2 | *MSH2\|NDUFAF2* |
| GO:0008283 | Cell population proliferation | 1 | *COL4A3* |
| GO:0042254 | Ribosome biogenesis | 1 | *P50914* |
| GO:0030198 | Extracellular matrix organization | 1 | *COL4A3* |
| GO:0044281 | Small molecule metabolic process | 1 | *ELOVL7* |
| GO:0022607 | Cellular component assembly | 1 | *NDUFAF2* |
| GO:0015031 | Protein transport | 1 | *P50914* |
| GO:0000278 | Mitotic cell cycle | 1 | *MSH2* |
| GO:0050877 | Nervous system process | 1 | *COL4A3* |
| GO:0051186 | Cofactor metabolic process | 1 | *ELOVL7* |
| GO:0055085 | Transmembrane transport | 1 | *KCNK12* |
| GO:0065003 | Protein-containing complex assembly | 1 | *NDUFAF2* |
| GO:0034655 | nucleobase-containing compound catabolic process | 1 | *P50914* |
| GO:0009790 | embryo development | 1 | *MSH2* |
| GO:0006605 | protein targeting | 1 | *P50914* |
| GO:0003013 | circulatory system process | 1 | *COL4A3* |
| GO:0006629 | lipid metabolic process | 1 | *ELOVL7* |
| GO:0006464 | cellular protein modification process | 1 | *ERCC8* |
| GO:0006790 | sulfur compound metabolic process | 1 | *ELOVL7* |
| GO:0006412 | translation | 1 | *P50914* |
| GO:0007005 | mitochondrion organization | 1 | *NDUFAF2* |

**Table S4** Mean kinship across all sites and within site mean kinship values. Based on simulations using allelic frequences, mean kinship values represent the order of relationships as follows: 0.059 are first cousins, 0.118 are half-siblings, and 0.236 are full sibling/parent-offspring relationships

| Site | All site MK (± SE) | Within site MK (± SE) |
| --- | --- | --- |
| Richmond Range | 0.098 (0.002) | 0.014 (0.002) |
| Port Macquarie  *Limeburners Creek*  *Lake Innes* | 0.088 (0.001)  *0.135 (0.002)*  *0.108 (0.001)* | 0.019 (0.001)  0.010 (0.001)  0.013 (0.001) |
| Kanangra-Boyd | 0.163 (0.001) | 0.018 (0.001) |
| Woronora Plateau  *Georges River*  *Upper Nepean* | 0.118 (0.002)  *0.187 (0.002)*  *0.138 (0.002)* | 0.030 (0.001)  0.017 (0.002)  0.009 (0.002) |
| Southern Tablelands | 0.127 (0.001) | 0.017 (0.001) |
| Narrandera  *East*  *West* | 0.109 (0.001)  *0.109 (0.002)*  *0.118 (0.002)* | 0.013 (0.001)  0.011 (0.001)  0.008 (0.002) |

**Table S5** Genomic positions of the 34 candidate loci identified via a Random Forest analysis as predictive of chlamydia infection in koalas including the scaffold, gene start (Start) and end (End) position, reference (Ref) and alternate (Alt) allele, location of loci (Func.refGene), gene name in annotation file (Gene.refGene), where relevant distance of loci from closest genes (GeneDetail.refGene), gene name(s) identified via a blastp search (Gene[s]), whether links to humans infected with *Chlamydia trachomatis* have been made (Links to *Chlamydia trachomatis*), and links to the publications. Attached as an Excel spreadsheet.

**Table S6** Gene Ontology (GO) terms and associated genes from Random Forest analysis of candidate loci in intronic regions predictive of chlamydia infection in koalas

| GO term ID | GO term definition | No. of genes | Genes |
| --- | --- | --- | --- |
| GO:0007049 | Cell cycle | 3 | *INCENP\|MLH3\|RASSF2* |
| GO:0022607 | Cellular component assembly | 2 | *MLH3\|TLN2* |
| GO:0051276 | Chromosome organization | 2 | *INCENP\|MLH3* |
| GO:0007059 | Chromosome segregation | 2 | *INCENP\|MLH3* |
| GO:0006464 | Cellular protein modification process | 2 | *RASSF2\|RNF11* |
| GO:0000278 | Mitotic cell cycle | 1 | *INCENP* |
| GO:0007155 | Cell adhesion | 1 | *TLN2* |
| GO:0007165 | Signal transduction | 1 | *RASSF2* |
| GO:0009056 | Catabolic process | 1 | *RNF11* |
| GO:0000003 | Reproduction | 1 | *MLH3* |
| GO:0006950 | Response to stress | 1 | *MLH3* |
| GO:0034330 | Cell junction organization | 1 | *TLN2* |
| GO:0034641 | Cellular nitrogen compound metabolic process | 1 | *MLH3* |
| GO:0042592 | Homeostatic process | 1 | *RASSF2* |
| GO:0048856 | Anatomical structure development | 1 | *RASSF2* |
| GO:0051301 | Cell division | 1 | *INCENP* |
| GO:0007010 | Cytoskeleton organization | 1 | *TLN2* |
| GO:0140014 | Mitotic nuclear division | 1 | *INCENP* |
| GO:0006259 | DdNA metabolic process | 1 | *MLH3* |

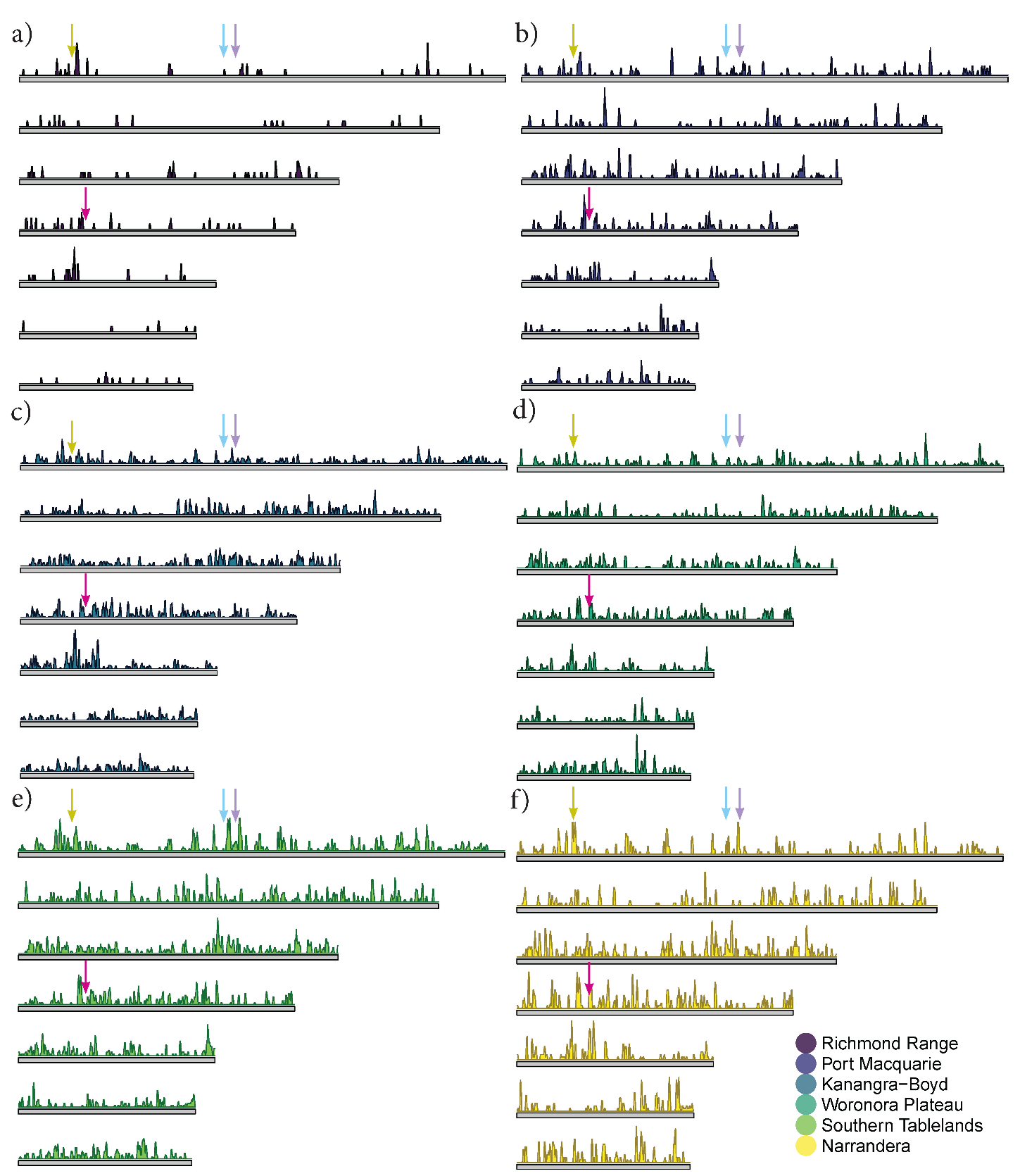

**Figure S1** Runs of homozygosity (>1,000 kb) in koalas across New South Wales Sentinel sites of a) Richmond Range; b) Port Macquarie, c) Kanangra-Boyd, d) Woronora Plateau, e) Southern Tablelands, and e) Narrandera across the seven longest scaffolds from the koala genome representing the autosomes. The height of the peak shows the density of the ROH across the scaffolds, higher peaks represent more ROH in a region. The arrows represent ROH present only in chlamydia positive individuals (*N* > 17 koalas) encompassing the following genes of un-annotated (yellow arrow), COL4A3 (blue arrow), ZSWIM6, SMIM15, NDUFAF2, ERCC8, RPL14, ELVL7, DEPDC1B (purple arrow), and EpCAM, MSH2, KCNK12, MSH6 (pink arrow). Gene Ontology pathways for these genes presented in Table S3.

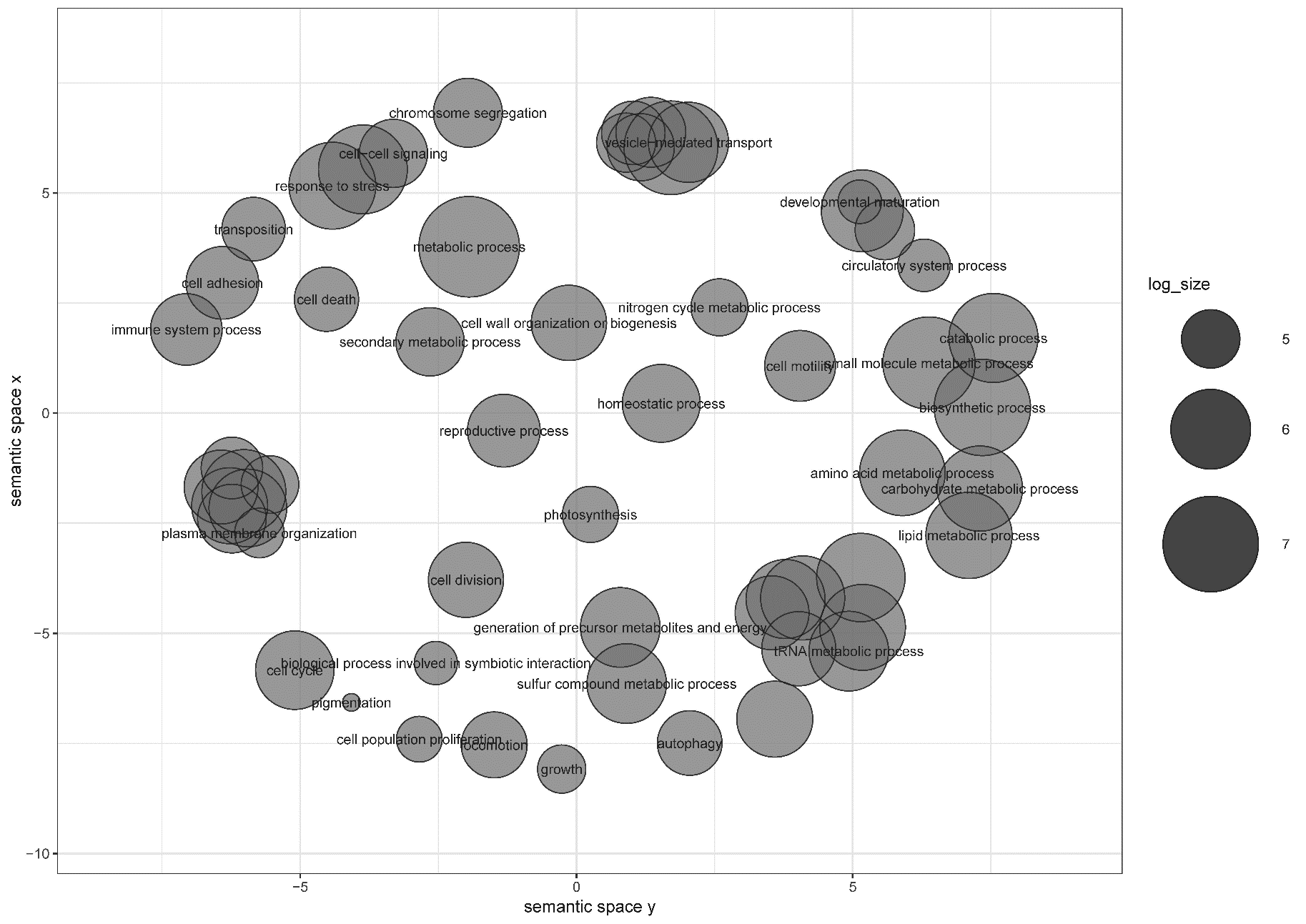

**Figure S2** Scatterplot showing clusters representative of enriched biological process Gene Ontology (GO) terms for the genes with run of homozygosity in chlamydia positive koalas only.

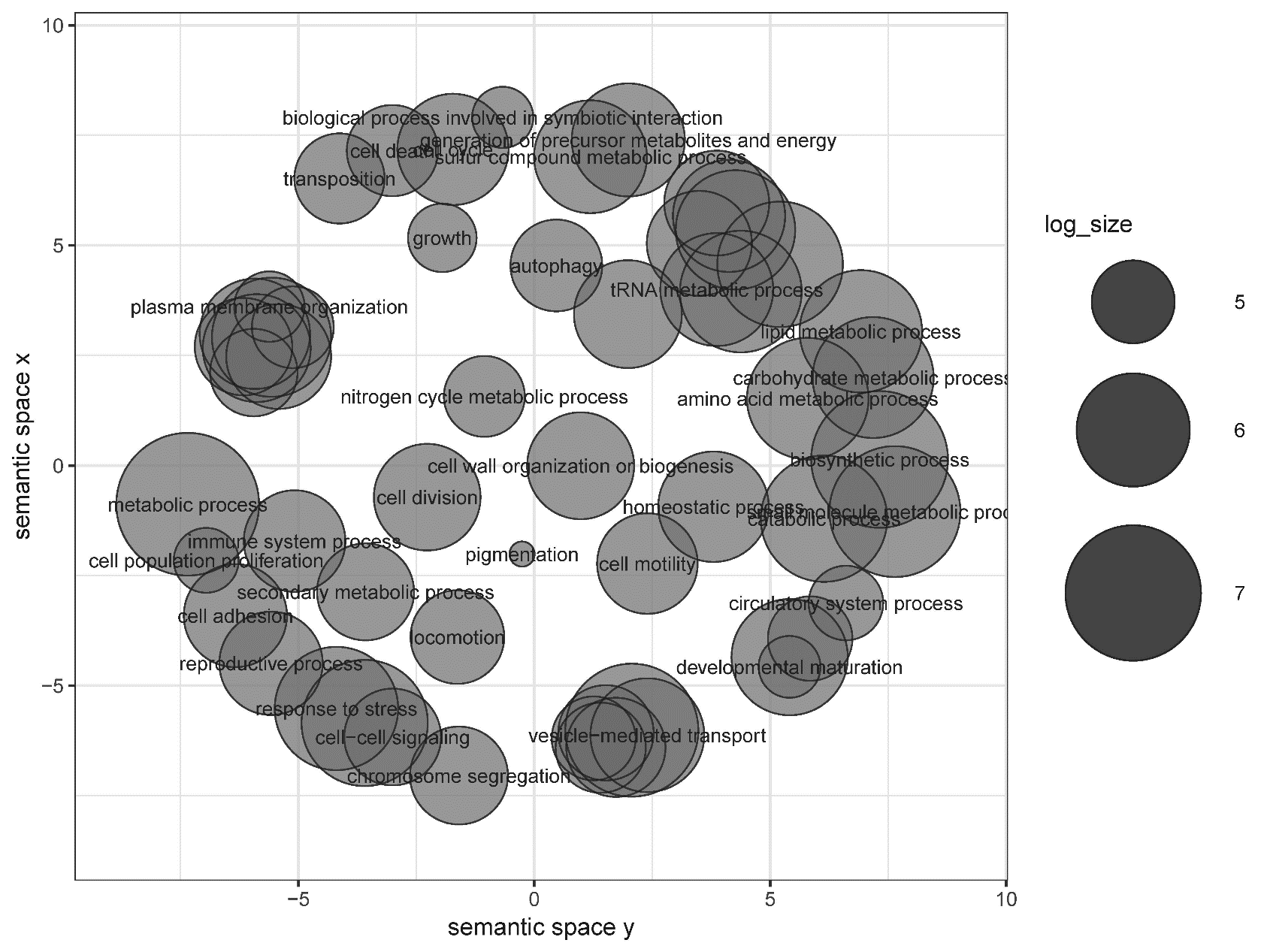

**Figure S3** Scatterplot showing clusters representative of enriched biological process Gene Ontology (GO) terms for the genes associated with the 34 candidate loci associated with chlamydia infection in koalas.
